# Disentangling the stomatal response to transpiration and relative humidity

**DOI:** 10.1101/2025.06.03.657603

**Authors:** Bar Ben Zeev, Or Shapira, Adi Yaaran, Yotam Zait

**Author notes:** Corresponding Autor: Yotam Zait.

## Abstract

Stomatal responses to atmospheric conditions are critical for balancing plant CO2 gain with water loss. Yet, the question of whether stomata respond directly to humidity or to the rate of water loss (transpiration) remains unresolved due to the tight coupling between these variables. To decouple them, we manipulated water vapor diffusivity using Helox (21% O_2_, 79% He), where vapor diffuses 2.33 times faster than in air, and Argox (21% O_2_, 79% Ar), where it diffuses 0.75 times slower. This allows control of VPD and transpiration independent of one another while maintaining a constant leaf temperature. We investigated the stomatal conductance (gsw) responses to air and Helox in several plant species, including Solanum lycopersicum, Vitis vinifera, and the fern Nephrolepis exaltata. In addition, we tested the Commelina communis, with coupled gas exchange and real-time imaging of stomatal aperture in intact leaves. In all angiosperms tested, we observed that stomatal closing was triggered exclusively by reductions in RH rather than by transpiration increases. When transpiration increased, but relative humidity remained constant, we detected an increase in gsw without a following decrease in the experimental time span. Exposure to Argox without altering RH reduced transpiration, likely enabling restoration of epidermal turgor and inducing passive mechanical closure. However, the rate of stomatal closure under reduced RH was 2.35 times faster than that observed under Argox, suggesting that passive turgor-driven responses cannot fully explain the kinetics or extent of closure under dry air. This work provides direct experimental evidence for humidity-driven stomatal closure.

## Introduction

Stomata are microscopic pores on the leaf surface that regulate gas exchange between plants and the atmosphere, balancing carbon dioxide uptake for photosynthesis with water loss through transpiration (Darwin, 1898). Stomata respond dynamically to environmental conditions; therefore, understanding the cues that trigger their opening and closure is central to predicting plant water use and stress responses. Vapor pressure deficit (VPD), defined as the difference in water vapor concentration between the leaf interior and the surrounding air, is a key determinant of transpiration rate and is widely considered a major driver of stomatal closure under dry atmospheric conditions (Buckley, 2005). Long exposure to high VPD commonly triggers stomata closure in angiosperms to avoid excessive water loss (Raschke, 1975). However, because VPD and transpiration (*E*) are inherently linked, it remains unresolved whether stomata respond to the driving force (VPD) itself or to the resulting water flux (*E*) or its consequences such as changes in osmolarity, turgor, water potential, or water content.

The debate over whether stomata respond to VPD or transpiration has persisted for decades. Previous efforts to isolate the environmental cues triggering stomatal closure yielded conflicting interpretations. A key contribution came from Mott and Parkhurst (1991), who developed an elegant method to decouple transpiration from relative humidity by using Helox, a gas mixture in which water vapor diffuses more than twice as fast as in air (Egorov and Karpushkin, 1988). By maintaining constant RH while increasing vapor diffusivity, they uncoupled the inherent link between RH and *E*, and observed that stomatal conductance and aperture decreased as *E* increased. This led them to propose that stomata respond directly to water loss itself rather than to RH or VPD. Their results supported the idea of localized “water loss sensors” within the leaf.

Nevertheless, other studies highlight humidity itself as a direct cue for stomata regulation, independent of water loss. This is supported by evidence of guard cells response to the RH gradients of the air layer surrounding the stomatal pore (peristomatal region) (Lange et al., 1971; Lösch, 1977; Appleby and Davies, 1983; Shope et al., 2008) or response to water loss via cuticular transpiration (Frensch and Schulze, 1988; Meinzer et al., 1997). For instance, Lang et al., (1970) demonstrated stomatal closure on isolated epidermis exposed to low RH from one side and constant high RH from other side and suggested that guard cells can function as “humidity sensors”controlled by their individual transpiration conditions (peristomatal transpiration). Later, Shope et al. (2008) used a similar set up, and showed that stomatal apertures of *T. pallida* decreased when VPD was increased, regardless of whether the epidermis contained live or dead epidermal cells, suggested that epidermal cells are not required for closure under high VPD.

Here we extended the approach of Mott and Parkhurst (1991) by introducing Argox (21% O_2_, 79% Ar), a mixture with reduced diffusivity, alongside Helox (21% O_2_, 79% He), enabling bidirectional control of transpiration while keeping RH constant. Notably, we coupled this setup with real-time microscopic imaging of stomata in intact leaves, allowing simultaneous quantification of stomatal aperture and gas exchange. This experimental setup enables us to revisit the classic question of whether stomata respond to the driving force (VPD) or the resulting flux (*E*) with higher spatial and temporal resolution and to examine this across multiple plant species.

## Material and methods

### Plant material

The study was conducted in a temperature-controlled greenhouse, with day/night temperatures maintained at 28/22°C. Four plant species were used: *Commelina communis* (Commelina), *Vitis vinifera* (Grapevine), *Solanum lycopersicum* (Tomato) and *Nephrolepis exaltata* (Boston fern). Plants were grown in 10 L pots filled with a growth medium composed of tuff, coconut, and peat in varying proportions (Tuff Marom-Golan, Ram 6, Israel). The plants were irrigated to container capacity twice a day, to ensure complete soil saturation. The irrigation solution contained a 6-6-6 NPK liquid fertilizer (ICL, Shefer Super, Israel) at a concentration of 0.5 mL/L. Measurements were taken on the youngest fully expanded leaf of each plant.

### Gas exchange measurements

Gas exchange measurements were conducted using the LI-6800F portable photosynthesis measurement system (LiCor Inc., Lincoln, NE, USA). Chamber type and size were adapted to the morphology of each species. For the *Solanum lycopersicum* (Tomato) and *Vitis vinifera* (Grapevine), a 6 cm^2^ was used. For *Nephrolepis exaltata* (Boston fern) a 2 cm^2^ fluorescence leaf chamber (6800-01A) was used, and in the *Commelina communis*, a clear top chamber (6800-12A), with leaf area of 2×3 cm was used; In this setup, the standard plastic cover was replaced with a 2 mm thick transparent glass cover to improve optical properties.

We used a 100 W Xenon light source Photonic F1 led (PHOTONIC Optische Geräte GesmbH & Co KG, Vienna, Austria) for supplementary light of 800 μmol photon m^−2^ s^−1^. The flow rate was set to 700±10 μmol s^-1^, and the fan speed was set to 5000 rpm. The reference CO_2_ concentration level was set at 420 μmol CO_2_ mol^−1^ air. The humidity in the chamber was regulated using the H_2_O reference controller. Light intensity was set at saturation intensity for each plant type, based on their respective light response curves; for *Solanum lycopersicum* 1000 μmol photon m^−2^ s^−1^, *Vitis vinifera* 1200 μmol photon m^−2^ s^−1,^ and for *Nephrolepis exaltata* 800 μmol photon m^−2^ s^−1^. The chamber air temperature was set at 25°C.

Measurements were conducted over a 2-hour period for each plant, with data logged at one-minute intervals (n=3). Infrared gas analyzers for the reference and sample were matched every 30 minutes to minimize drift. It is important to note that the vapor pressure of water is a function of temperature alone and remains constant regardless of the carrier gas (e.g., air, Helox, or Argox), as described by Dalton’s law of partial pressures (Wiegleb, 2023). To calculate the ratios between the stomatal conductance (*g*_*sw*_) or transpiration rate (*E*) in the different gas mixtures (air, Helox, and Argox) or relative humidity, we averaged the values recorded during the intervals of 15 - 30 minutes and 60 - 90 minutes in each experiment.

### IRGA calibration for gas mixtures

Because the current LI-COR 6800 infra-red gas analyzer (IRGA) system is factory-calibrated for measurements in ambient air, it cannot directly provide accurate gas exchange measurements in non-standard gas mixtures such as pure N_2_/O_2_ blends. This is because the IRGA assumes specific physical properties of air, like the density and size of the molecules, that change with gas composition. These differences can lead to incorrect readings of gas concentrations. To calibrate the IRGA for measurements under different gas mixtures (Helox or Argox), a calibration curve was generated using an empty leaf chamber (Fig.S1,S2). For water vapor calibration (H_2_O), the “Humidifier” and the “Desiccant” controls were manually adjusted to create a range of humidity levels. Once stable readings were obtained under ambient air, the gas mixture was switched to Helox or Argox, and measurements resumed after the system stabilized. All H_2_O calibrations were conducted with the CO_2_ scrubber turned off to avoid excessive desiccation by the soda lime scrub and maintain ambient CO_2_ concentrations.

The CO_2_ calibration was performed by manually adjusting the CO_2_ injector to achieve a range of concentrations under both ambient air and gas mixtures. During this procedure, the H_2_O controller was turned off, and the CO_2_ scrubber was kept activated constantly to avoid background corrections of the CO_2_ concentration by the system. All calibrations were conducted at a constant air temperature of 25°C. After calibration, we used the equations provided by the LI-COR-6800 with the calibrated values.

It is important to note that all measurements were conducted at a constant temperature to avoid confounding effects on the diffusion coefficient and to exclude the direct influence of temperature on stomatal conductance (Mills et al., 2024).

### Boundary layer conductance calibration

The boundary layer conductance (*g*_*bl*_) also depends on the diffusion coefficient of the gas mixture, which varies with different gas mixtures. Therefore, *g*_*bl*_ was calibrated separately for each gas mixture used in our experiments (air, Helox, Argox). The boundary layer conductance (*g*_*bl*_) was calibrated for each gas mixture (air, Helox, Argox). To determine the boundary layer thickness, we used the empirical relationship:

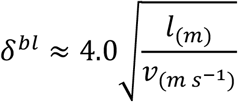

Where *l* is the leaf length in the direction of the airflow (in meters), *v* is a symbol for ambient wind velocity, in m s^-1^. *δ*^*bl*^ represent the thickness of the boundary layer, in millimetres. The coefficient 4.0 is use as a factor for the air temperature in the boundary layer development at 20°C-25°C (Nobel, 2020). In our experiment, the wind velocity was set to 2 m s^-1^ following (Shapira et al., 2024). The characteristic length *l* was taken as the chamber diameter, as we ensured that the leaf occupied the chamber entirely in the direction of flow. To convert the estimated thickness of the boundary layer to boundary layer conductance for each gas mixture, we used the following equation (Aphalo and Jarvis, 1993):

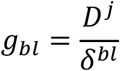

Where *D*^*j*^ is the diffusion coefficient of water vapor in the background mixture *j* (e.g., H_2_O or CO_2_ in air, Helox, or Argox). The diffusion coefficient of water vapor in air and Helox were taken from (Parkhurst and Mott, 1990) and of Argox from (O’Connell et al., 1969).

### Artificial leaf validation

We used an artificial leaf made of two water-resistant plastic disks to validate the experimental system and boundary layer conductance estimates. The abaxial (lower) disk contained pores with a diameter of 500 µm in a density of 100 pores cm^-2^, the adaxial disk contained pores with a diameter of 500 µm in a density of 10 pores cm^-2^. A piece of filter paper (Whatman No. 3) saturated with distilled water was placed between the plastic disks to serve as a water source. To ensure continuous hydration, the tip of the filter paper was immersed in a beaker filled with distilled water. The artificial leaf was inserted into a 6 cm^2^, fluorescence leaf chamber (6800-01A). As in all experiments, the air temperature within the chamber was maintained at 25°C.

### Stomatal imaging and aperture measurement

To monitor the stomatal aperture of the *Commelina communis*, we used a Stereo microscope (Nexcope NSZ-818, Nexcope, Ningbo, China), equipped with a 1× apochromatic plan objective and a magnification range of 0.75× to 13.5×. Imaging was performed using a 4K digital camera (TOP4KB, Topika, Israel).

The LI-COR 6800 gas exchange connected to a clear top chamber (6800-12A) was positioned directly under the microscope lens, and the scale was calibrated at the maximum magnification setting (13.5×) to ensure accurate dimensional analysis. Stomatal images were captured every 1 minute in synchrony with gas exchange data logging.

Image analysis was performed using ImageJ software (Rasband, W.S., ImageJ, US National Institutes of Health, Bethesda, MD, USA, http://imagej.nih.gov/ij/, 1997– 2025). For each time point, the same five stomata were tracked across the time series, and stomatal aperture was quantified by measuring the width of each stoma.

### Statistical and data analysis

Statistical analyses and data visualization were performed using R statistical software, version 4.4.3 (https://cran.r-project.org) with the ggplot2, dplyr, and ggpubr packages. Differences between treatment groups were assessed using two-sample t-tests. Statistical significance was defined as p < 0.05.

## Results

### Validation of the experimental system using an artificial leaf: disentangling the effects of VPD and gas diffusivity

To validate our experimental system and IRGA calibration, we tested an artificial leaf with fixed-size pores and constant stomatal conductance (*g*_*sw*_) in air (Fig 1). The transpiration rate (*E*) of the artificial leaf is variable and depends on the vapor pressure deficit (VPD). In the first experiment, we switched the background gas from air to Helox. As expected, E increased by a factor of 1.68, while g_sw_ increased by a factor of 2.17. Relative humidity (RH) remained approximately constant at 70% in both gases. In the second experiment, we reduced the RH from 70% to 54% in air. Under these conditions, the *g*_*sw*_ remained nearly unchanged, while the *E* is increased by a factor of 1.56 in response to the higher VPD, confirming that the artificial leaf responds to changes in VPD through *E*, but not via changes in conductance (Fig. 1).

**Figure 1:**
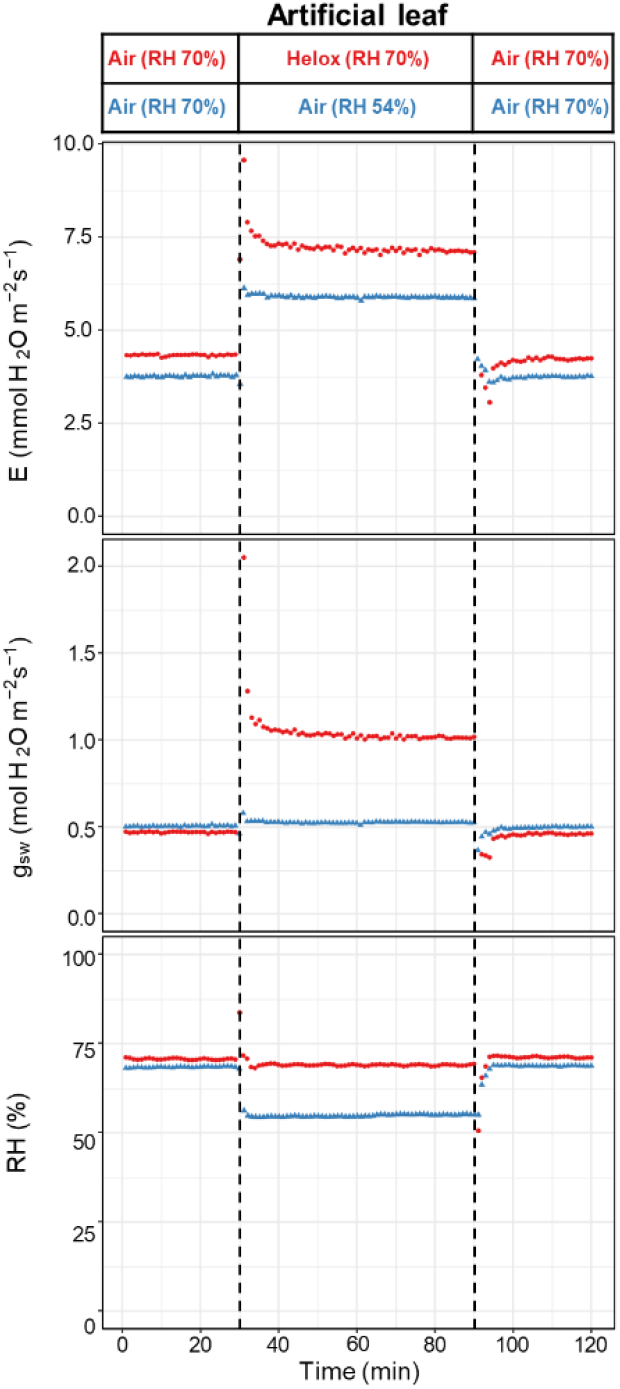
Artificial leaf shows constant conductance in response to changes in RH and variable conductance in response to Helox. Transpiration rate (*E*), stomatal conductance (*g*_*sw*_), and relative humidity (RH) over time during two artificial leaf experiments. In the “changing RH” experiment (blue triangles), RH was altered while the gas composition remained constant. In the “changing gas mixture” experiment (red circles), the gas was switched from air to Helox while maintaining constant RH. Dashed vertical lines mark the start and end of each treatment. The box above each panel shows the RH and gas mixture treatment in each experiment.

### Stomatal conductance response to transpiration rate and relative humidity in different species

To test whether stomata respond to the driving force for water loss (i.e., RH/VPD) or to the actual water flux (transpiration, *E*), we increased vapor diffusivity by using Helox while maintaining constant RH, thereby decoupling *E* from RH. First, we switched from air to Helox and then back to air. Next, we reduced RH from 75% to 54% and then restored it to 75%, inducing an equivalent to the Helox increase in *E*. We applied both treatments to three species representing different growth forms: *Solanum lycopersicum* (herbaceous), *Vitis vinifera* (woody), and *Nephrolepis exaltata* (pteridophyte).

In all the three species, switching to Helox increased *E* and *g*_*sw*_, while RH remained constant. In contrast, reducing RH produced a similar increase in *E*, but it was followed by a reduction in *E*. Also, it triggered a reduction in *g*_*sw*_, both indicating that stomatal closure was driven by RH rather than by water loss. In *Solanum lycopersicum*, Helox increased *E* by 1.78-fold and *g*_*sw*_ by 2.46-fold, while RH reduction increased *E* by 1.32-fold and decreased *g*_*sw*_ to 0.81 (Fig. 2A). In *Nephrolepis exaltata*, Helox increased *E* and *g*_*sw*_ by 1.71 and 1.92, respectively, whereas RH reduction caused only a small *E* increase (1.33-fold) and no change in *g*_*sw*_ (Fig. 2B). In *Vitis vinifera*, Helox increased both *E* and *g*_*sw*_ by 1.3-fold, while RH reduction increased E by 1.27-fold and slightly decreased *g*_*sw*_ to 0.84 (Fig. 2C).

**Figure 2:**
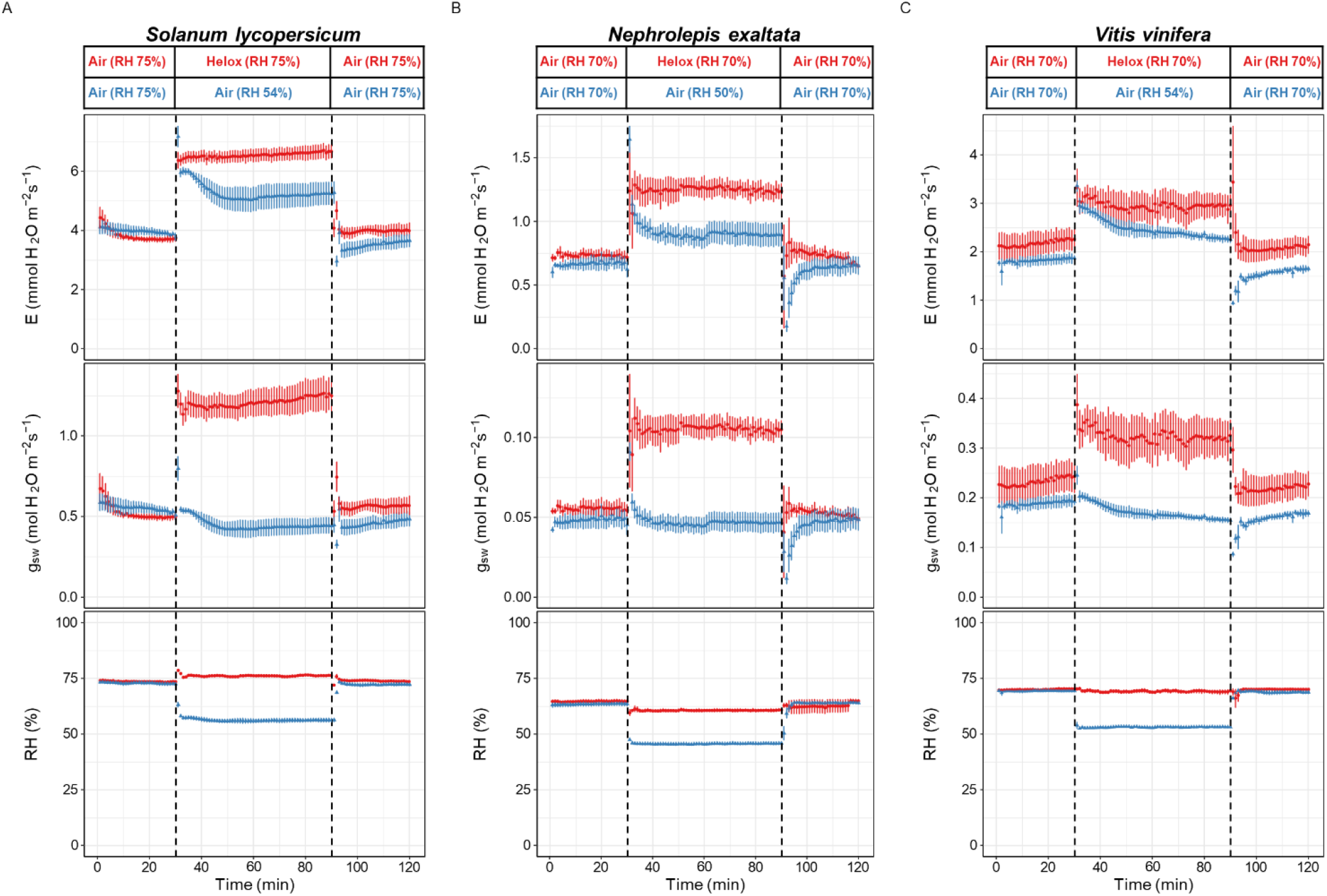
Stomatal conductance responds more strongly to reduced relative humidity than to increased transpiration across diverse plant species. Transpiration rate (*E*), stomatal conductance (*g*_*sw*_), and relative humidity (RH) over time (minutes) in *Solanum lycopersicum* (A), *Nephrolepis exaltata* (B), and *Vitis vinifera* (C), measured under two experimental conditions. Blue triangles indicate the “changing RH” experiment (75% to 54% and back to 75% RH), and red circles indicate the “changing gas mixture” experiment (air to Helox and back to air). Dashed vertical lines mark the start and end of each treatment. The box above each panel shows the RH and gas mixture treatment in each experiment. Values represent means ± standard error (*n* = 3).

To determine whether changes in stomatal conductance under Helox occur independently of transpiration, we conducted an experiment in which *E* was held nearly constant by adjusting RH in parallel with the gas mixture. RH was lowered to ∼50% in air and raised to 75% in Helox, resulting in comparable *E* rates across treatments. Despite similar *E* values, *g*_*sw*_ increased under Helox in all cases: by 2.51 in the artificial leaf (Fig. 3A), 2.58 in *S. lycopersicum* (Fig. 3B), 1.65 in *N. exaltata* (Fig. 3C), and 1.56 in *V. vinifera* (Fig. 3D). These results indicate that the increase in *g*_*sw*_ under Helox occurs even when water flux (*E*) is not elevated.

**Figure 3:**
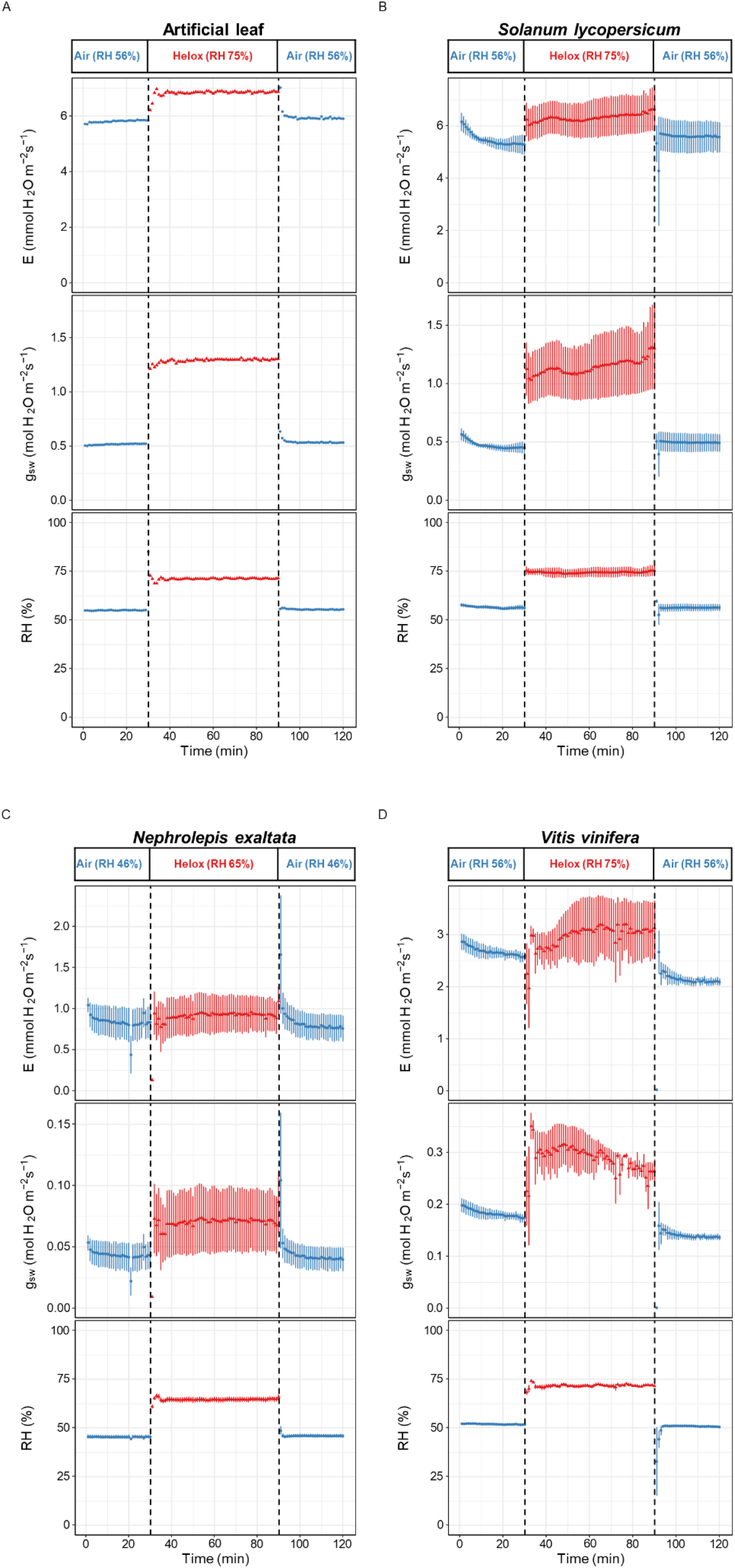
Increasing RH to compensate for high diffusivity in Helox maintained nearly constant transpiration and did not trigger stomatal closure. Transpiration rate (*E*), stomatal conductance (*g*_*sw*_), and relative humidity (RH) over time (minutes) in the artificial leaf (A) and three plant species: *Solanum lycopersicum* (B), *Nephrolepis exaltata* (C), and *Vitis vinifera* (D). Blue circles represent air with low RH; red triangles represent Helox with high RH. Dashed vertical lines mark the start and end of the treatment period. The box above each panel shows the RH and gas mixture treatment in each experiment. Values are means ± standard error (*n* = 3 for plants; *n* = 1 for artificial leaf).

### Simultaneous measurement of stomatal conductance and aperture in response to transpiration manipulation and relative humidity

To examine whether actual stomatal aperture regulation is compatible with gas exchange, and stomatal response to *E* and RH, we coupled gas exchange setup with real-time microscopic imaging of stomata in intact leaves of *Commelina communis*. Then, leaves were exposed to three treatments: Helox, reduced relative humidity, and Argox (Fig. 4). Argox, which has lower gas diffusivity than air, served as a counterpoint to Helox.

**Figure 4:**
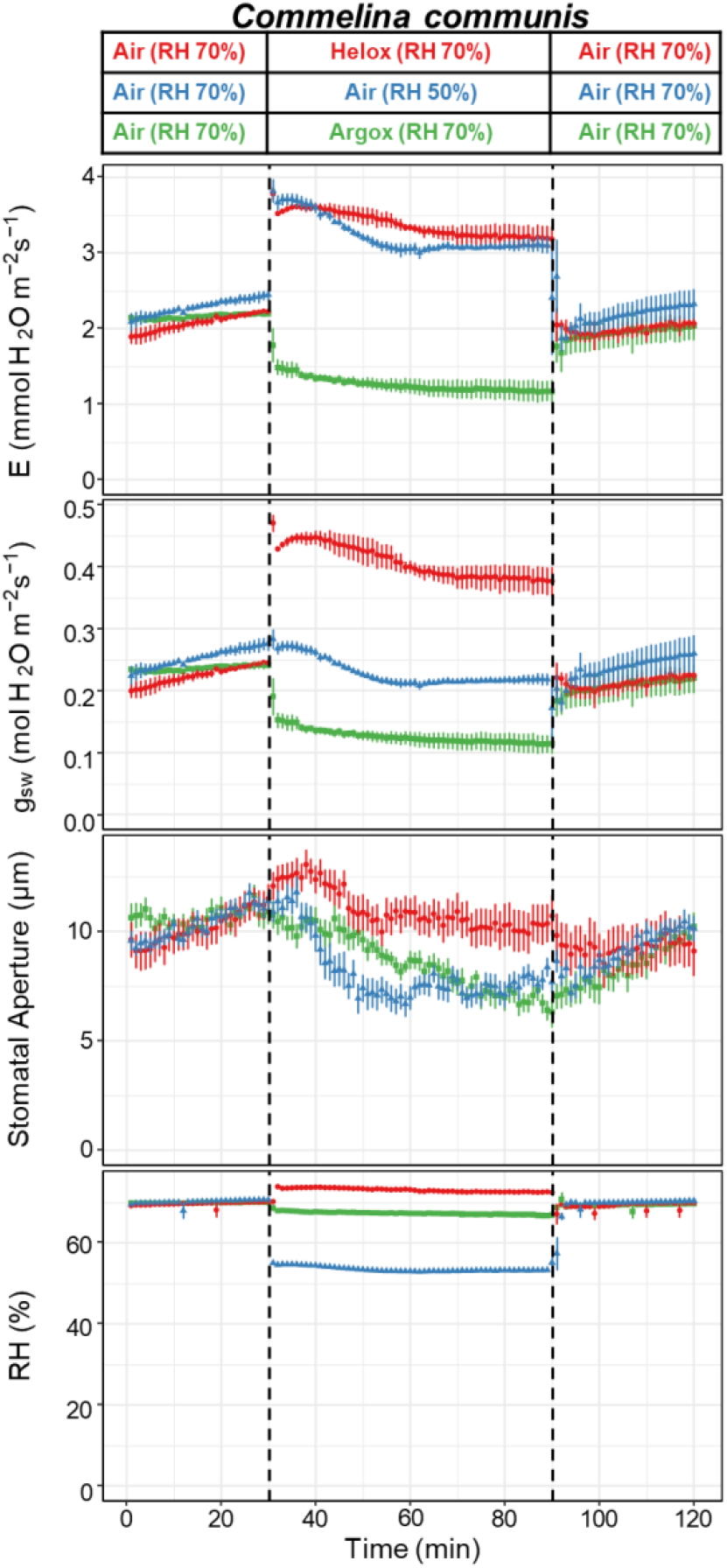
Stomatal aperture remains unchanged under high transpiration (Helox) but closes under low RH or Argox. Transpiration (*E*), stomatal conductance (*g*_*sw*_), stomatal aperture (pore width, µm), and relative humidity (RH) over time (minutes) in *Commelina communis* under three experimental conditions. Blue triangles represent the “changing RH” experiment (75% to 54% and back to 75% RH), red circles represent the “air to Helox” experiment (air to Helox and back to air), and green squares represent the “air to Argox” experiment (air to Argox and back to air). Dashed vertical lines mark the start and end of each treatment. The boxes above each panel show the fixed RH levels in the system. Values are means ± standard error (*n* = 3).

Switching the gas mixture from air to Helox increased *E* by 1.5 and *g*_*sw*_ by 1.62-fold, respectively. In contrast, the stomatal aperture exhibited a transient increase followed by a return to baseline, with no net change, indicating a decoupling between *g*_*sw*_ and aperture. On the other hand, switching the gas mixture from air to Argox reduced *E* by 46% and *g*_*sw*_ by 51%, with a corresponding decrease of 31% in the stomatal aperture.

Reducing RH from 70% to 50% on air increased *E*, similar to the increase observed in the switching from air to Helox. However, as opposed to the Helox effect, the humidity drop caused a reduction of 20% in *g*_*sw*_ and 31% in aperture. The stomatal closing rate in the RH reduction was 0.40 μm min^-1^ and only 0.17 μm min^-1^ in Argox. In Helox, there is approximately no change after the stomatal aperture returns to the baseline conductance. Together, these results show a stronger reduction in aperture following RH change and that *g*_*sw*_ can be modulated by transpiration rate independently of changes in stomatal aperture. Percent changes in *g*_*sw*_ and aperture across treatments are summarized in Figure 5.

**Figure 5:**
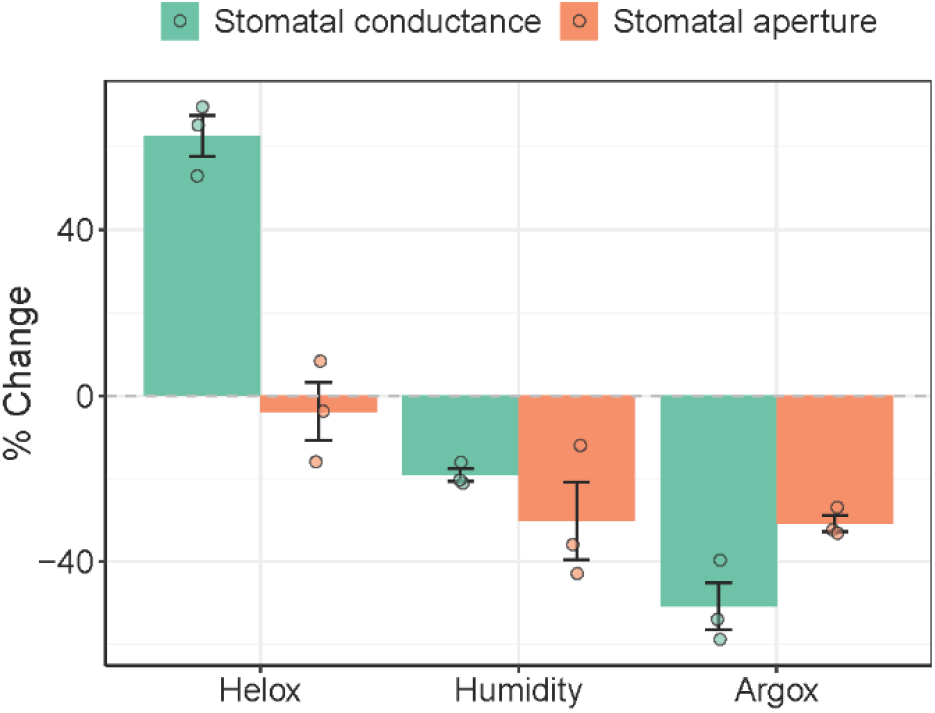
Stomatal conductance and aperture decrease under low RH and Argox but not under Helox. Percent change in stomatal conductance (turquoise) and stomatal aperture (orange) across three experimental treatments: Helox, low humidity, and Argox. Individual data points represent the average values recorded at T = 60–90 min (post-treatment) relative to T = 15–30 min (pre-treatment), calculated as: 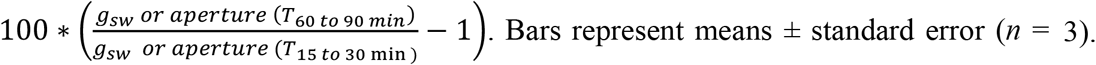.

## Discussion

Our results demonstrate that stomatal closure is triggered by a reduction in relative humidity (RH), not by elevated transpiration (*E*) per se (Figs. 1–4). This finding resolves a long-standing question in stomatal physiology: do stomata respond to the driving force for water loss (i.e., RH/VPD), or to the resulting water flux (*E*)?

In *Commelina communis*, when RH was held constant, Helox significantly increased *E* and *g*_*sw*_, yet the aperture remained unchanged (Fig. 4), demonstrating that guard cells can maintain pore opening even under elevated water flux if RH is not altered. In contrast, when RH was reduced, a consistent decline in aperture was observed, confirming that stomatal closure is triggered specifically by atmospheric dryness (Figs. 2, 4). Interestingly, the Boston fern, which lacks the ABA stomatal active response (Cardoso and McAdam, 2019), showed minimal reduction in *g*_*sw*_ following RH drop, supporting the idea that ABA takes part in an active humidity-sensing mechanism (Bunce, 1996).

Our findings are broadly consistent with previous Helox-based studies (Egorov and Karpushkin, 1988; Mott et al., 1991; Mott and Parkhurst, 1991), which reported increased *g*_*sw*_ under higher vapor diffusivity. However, Parkhurst and Mott also observed a reduction in stomatal aperture when *E* increased and RH remained constant, unlike the stable aperture we observed via direct imaging. This discrepancy may reflect differences in experimental design: (1) their lower baseline RH (53%) compared to our study (70%) could have influenced stomatal behavior due to altered water balance; and (2) aperture in their study was inferred from gas exchange, whereas we directly measured it using high-resolution microscopy. Additional support for our findings comes from (Shope et al., 2008), who observed similar stomatal apertures under air and Helox in *Vicia faba* epidermal peels when RH was constant. Likewise, Pieruschka et al., (2010) reported no difference in *E* or *g*_*sw*_ between air and Helox when radiation load (total energy absorbed by a leaf thru radiation) was controlled. Our experiments were also conducted under constant temperature and light intensity, reinforcing the conclusion that RH, not transpiration or energy balance, is the dominant signal driving stomatal closure under these conditions.

The use of Argox provided further insight into the mechanisms of closure. Exposure to Argox reduced *E* without altering RH, likely enabling restoration of epidermal turgor and inducing passive mechanical closure (Mott and Franks, 2001; Buckley et al., 2011; Zait et al., 2017). However, the rate of stomatal closure under low RH was 2.35 times faster than that observed under Argox (Fig. 5), suggesting that passive turgor-driven responses cannot fully explain the kinetics or extent of closure under dry air. This difference may allow us to distinguish between passive and active components of the stomatal response.

Under constant RH, the *g*_*sw*_ ratio between Helox and air nearly approached the theoretical value of 2.33 in the artificial leaf but varied across species (Fig. S3), possibly due to differences in stomatal anatomy, epidermal-guard cell mechanics (Pichaco et al., 2024), or stomatal size and density. Parkhurst et al. (1990) suggested that amphistomatous leaves, with shorter diffusion paths, may show higher *g*_*sw*_ ratios under Helox. However, in our study, we could not assess these anatomical or structural traits, and their contribution to the observed variation remains to be investigated.

To mechanistically dissect RH versus *E-*driven responses, we propose that each initiate distinct physiological pathway (Fig. 6). In the RH-driven scenario, *E* remains constant, and thus water status parameters, such as turgor pressure, osmotic potential, and relative water content, should also remain stable (Fig. 6A). Nonetheless, the stomatal aperture is reduced, implying an active sensing and signal transduction mechanism triggered by RH (Fig. 6B). This pathway likely involves both ABA-dependent and ABA-independent components, for example via the protein kinase OST1 (Yoshida et al., 2006; Merilo et al., 2018; Tulva et al., 2023) which may play a central role in RH-induced stomatal closure. In contrast, under Helox, *E* increases and may impact leaf water status, but this alone is insufficient to trigger closure, further supporting the idea that transpiration is not the direct stimulus (Fig. 6C).

**Figure 6:**
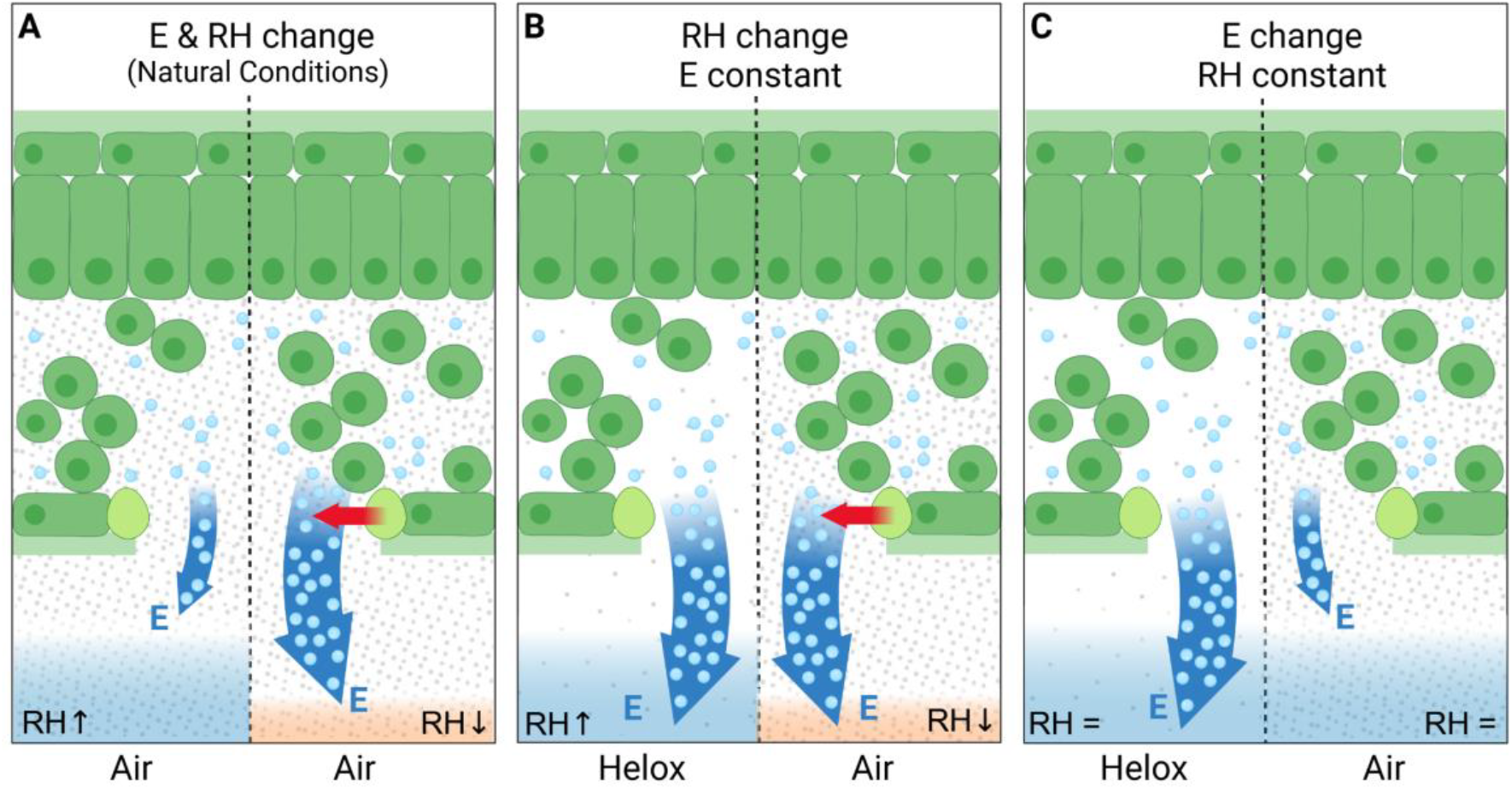
Stomatal closure is triggered by changes in relative humidity (RH), not by changes in transpiration rate (*E*). (A) Under natural conditions, a drop in RH (orange vs. blue background) increases the water vapor pressure gradient, leading to higher *E* (blue arrow) and stomatal closure (red arrow). (B) When the surrounding air (dense gray dots) is replaced with the higher diffusive Helox (sparse gray dots) while RH remains constant (blue background), *E* increases (blue arrow), but stomata remain open. (C) When the surrounding Helox (sparse gray dots) is replaced with the lower diffusive air (dense gray dots) while RH is proportionally increased (orange vs. blue background) to maintain constant *E* (blue arrow), stomatal closure still occurs (red arrow).

It is possible that the decline in RH within the substomatal airspace acts as a vapor-phase signal, sensed either directly by guard cells or through yet-unidentified leaf structures (Sibbernsen and Mott, 2010; Buckley, 2019). Recent hypotheses suggest that sub-saturation of the leaf airspace could be critical for initiating the stomatal closure in response to high VPD (Márquez et al., 2024), potentially via water potential gradients within the leaf suggesting that the stomata is sensing RH in the leaf interior.

Exogenous humidity sense mechanisms are unknown in land plants (Embryophytes) but serve as an important factor in other life forms like insects and arachnids (Merrick and Filingeri, 2019). While the RH sensor’s identity and location in plants remain unknown, our findings lay the groundwork for identifying the molecular and biophysical mechanisms by which plants sense and respond to atmospheric humidity.

## Acknowledgments and Funding

This work was supported by the Israel Science Foundation (ISF 2076/23) to **Y.Z**.

## Author Contributions

**B.B.Z., O.S., A.Y., and Y.Z**. jointly conceived the research and designed the experimental plan. **B.B.Z**. conducted the experiments and analyzed the data. **B.B.Z A.Y, and Y.Z**. wrote the manuscript.

## Supporting information

**Figure S1:**
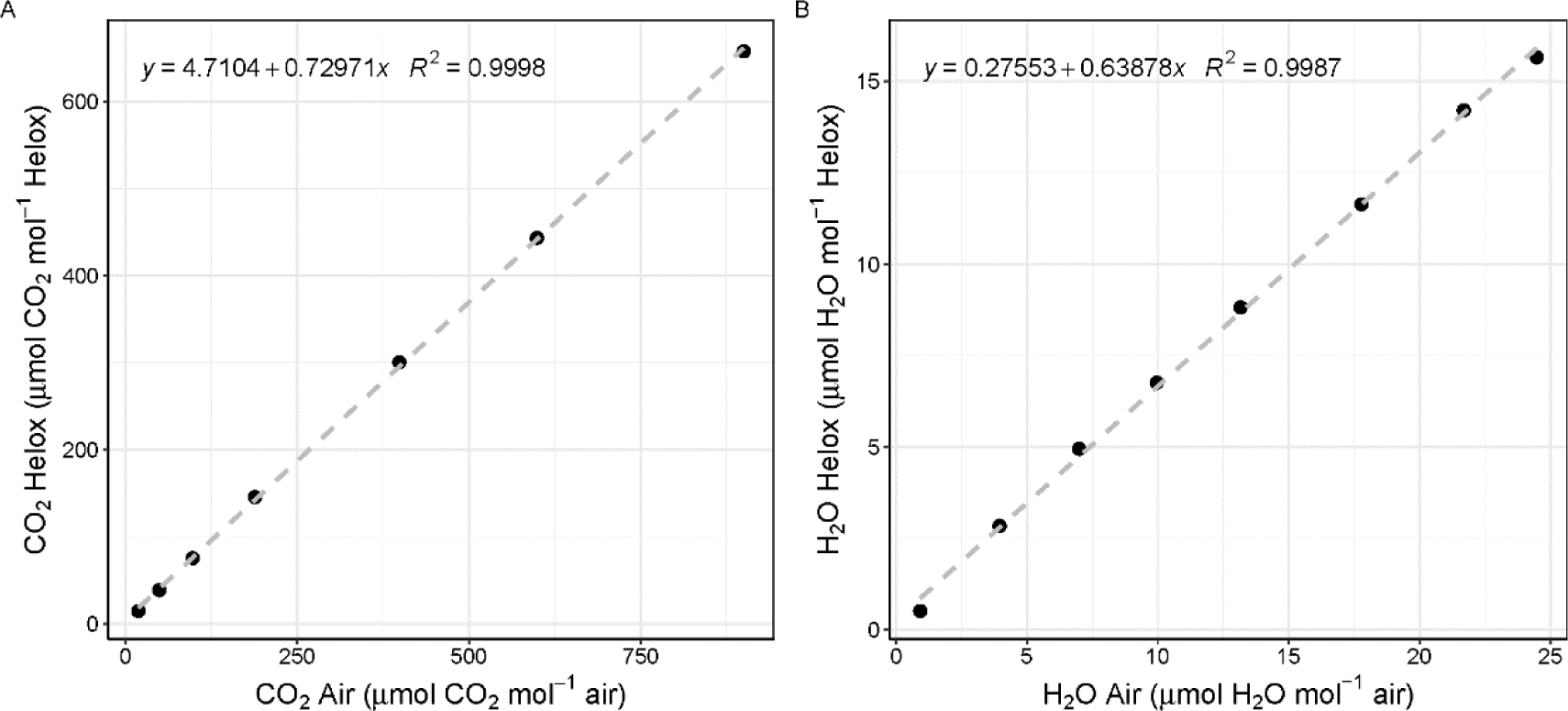
IRGA calibration to Helox mixture (21% O_2_, 79% He). (A) calibration curve of CO_2_ concentration in Helox and CO2 concentration in the air. (B) calibration curve of H_2_O concentration in Helox and H_2_O concentration in the air.

**Figure S2:**
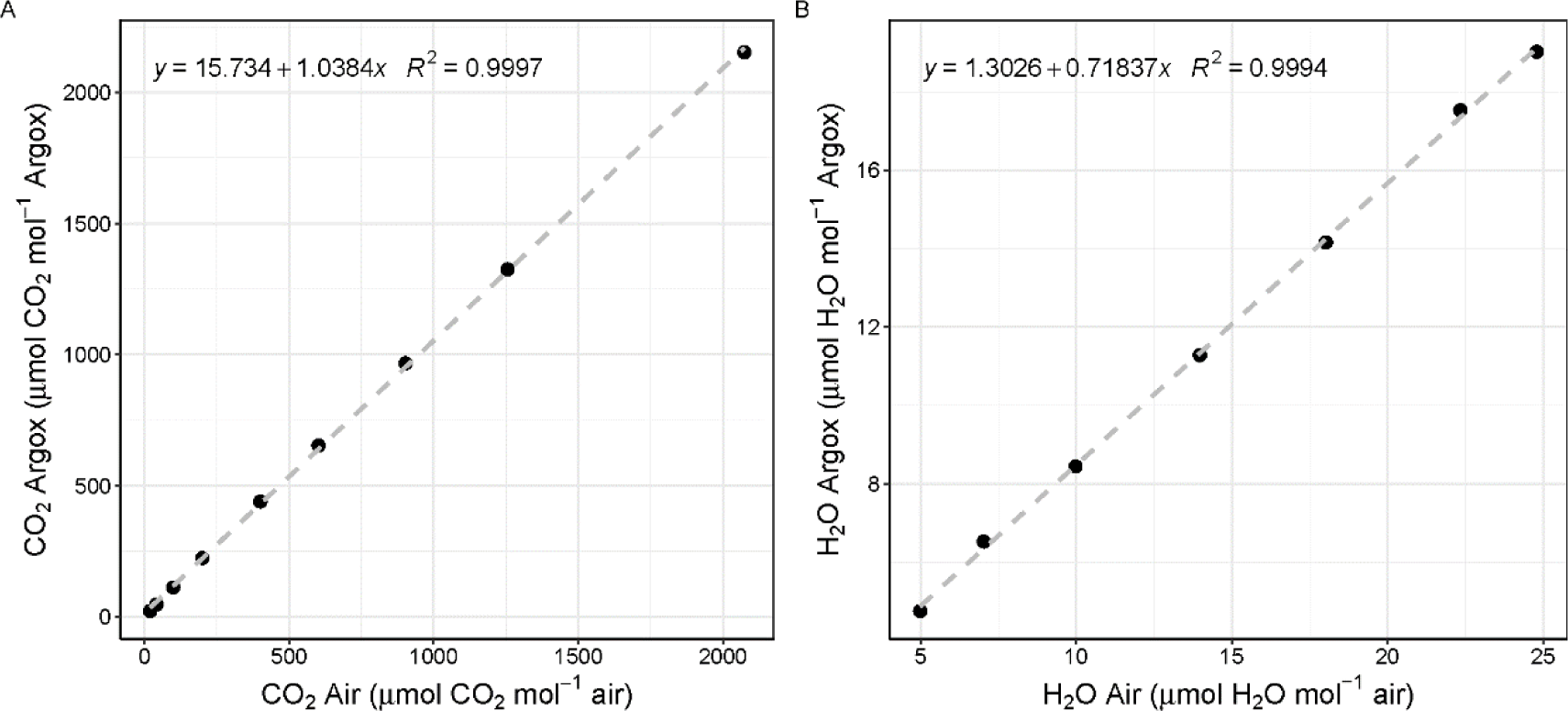
IRGA calibration to Argox mixture (21% O_2_, 79% Ar). (A) calibration curve of CO_2_ concentration in Argox and CO_2_ concentration in the air. (B) calibration curve of H_2_O concentration in Argox and H_2_O concentration in the air.

**Figure S3:**
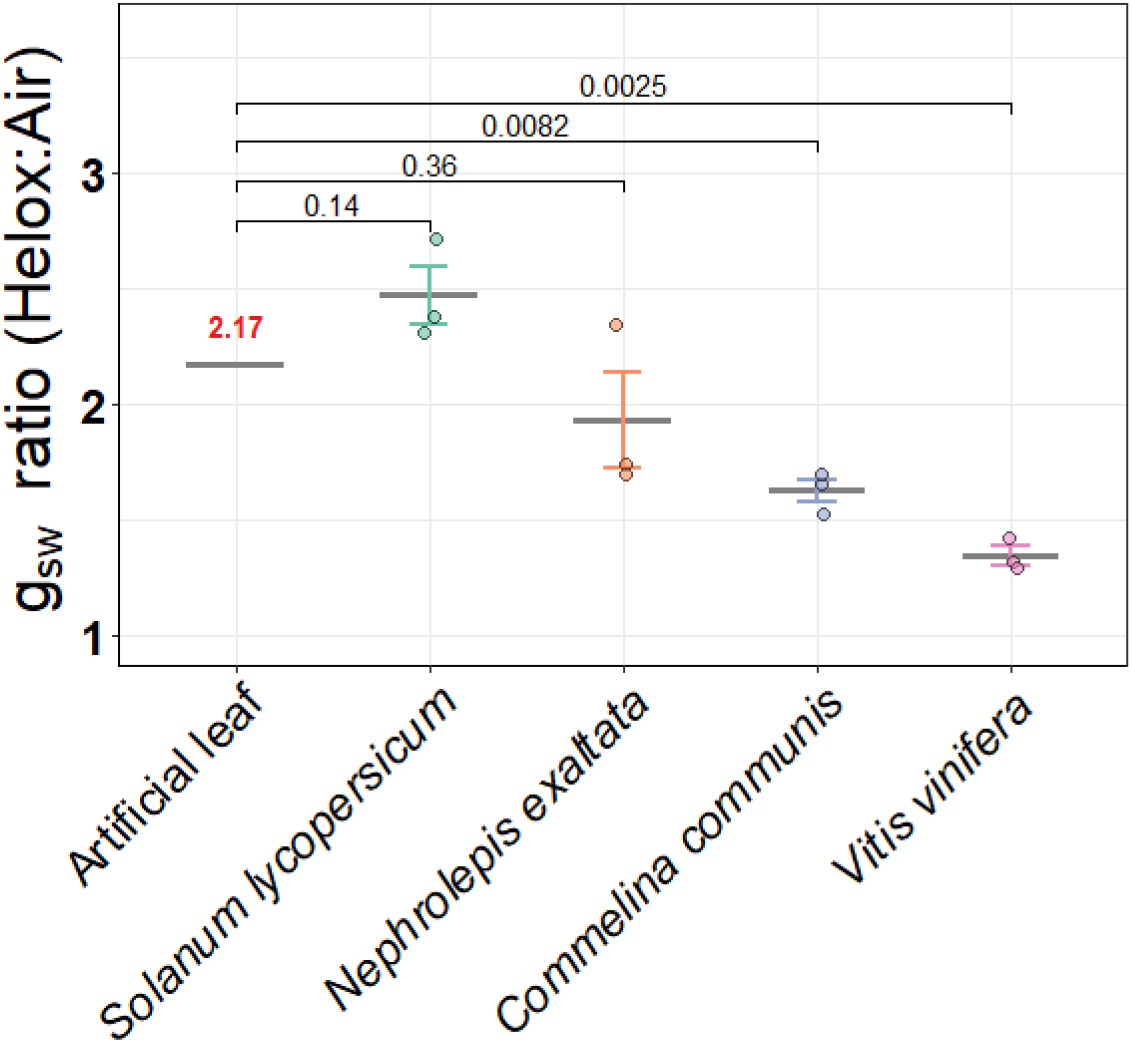
The ratio of stomatal conductance in Helox versus air varies among species. Ratio of stomatal conductance (*g*_*sw*_) measured in air versus Helox for the artificial leaf and four plant species. Gray horizontal line is the mean *g*_*sw*_ ratio, Individual data points are shown, and bars represent means ± standard error (*n* = 3) of values recorded between minutes 15 - 30 and 60 - 90 in each experiment.

